## Supplementary material for "The TFIIH Complex is Required to Establish and Maintain Mitotic Chromosome Structure": Supplmental_Figures

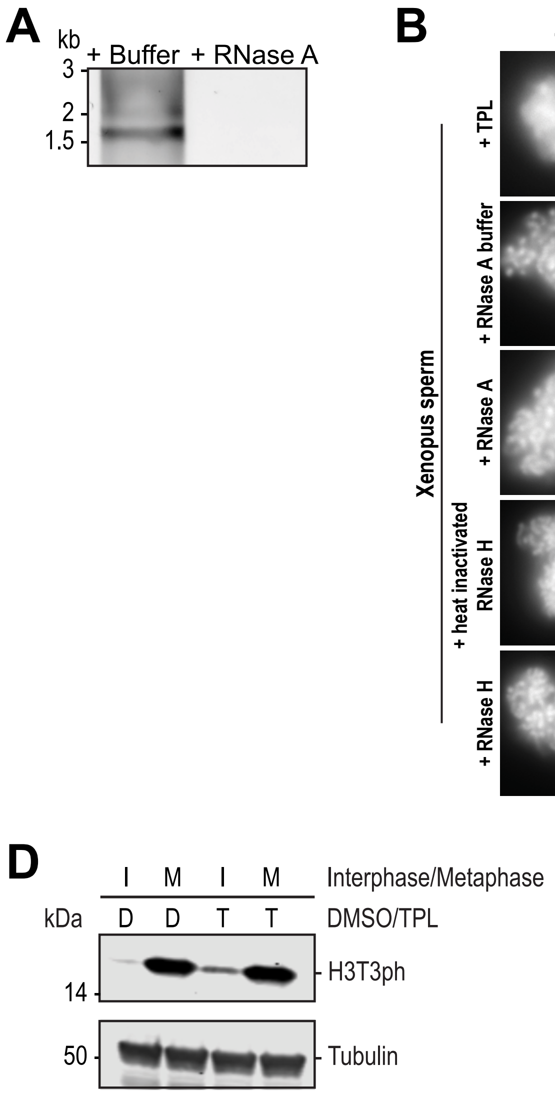


**Figure S1: Effects of RNases and various transcription inhibitors on chromosome condensation and mitotic progression:** **(A)** 1% agarose gel stained with SYBR safe stain showing RNase A dependent removal of total RNAs from high-speed supernatant egg extracts. **(B)** Representative fluorescence images of chromatid assembly at steady state (180 mins after sperm addition) in the presence of indicated inhibitors, buffers, and enzymes. See methods for details. **(C)** Representative fluorescence images of chromatid assembly at steady state (180 mins after sperm addition) in the presence of DMSO or 30 μM CDK7 inhibitor THZ1. **(D)** Triptolide does not affect M phase entry in egg extracts. Western blot for histone H3T3ph and Tubulin in extracts cycled from interphase to metaphase in the presence of DMSO or triptolide. H3T3ph is only highly phosphorylated in M phase, and its appearance is not affected by triptolide treatment.

**A**

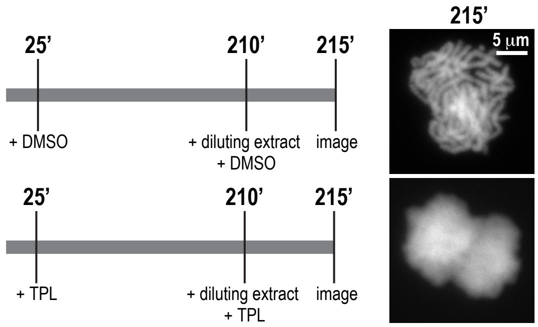


**Figure S2: Continuous TFIIH activity is required to maintain chromosome structure: (A)** Representative fluorescence images of *Xenopus* sperm nuclei incubated in extracts at indicated timepoints. DMSO or Triptolide were added 25 mins after addition of nuclei, and extracts were diluted 10-fold with extract containing DMSO or Triptolide at 210 mins after nuclei addition, and further incubated for 25 min before imaging. Chromatin was stained with Hoechst.

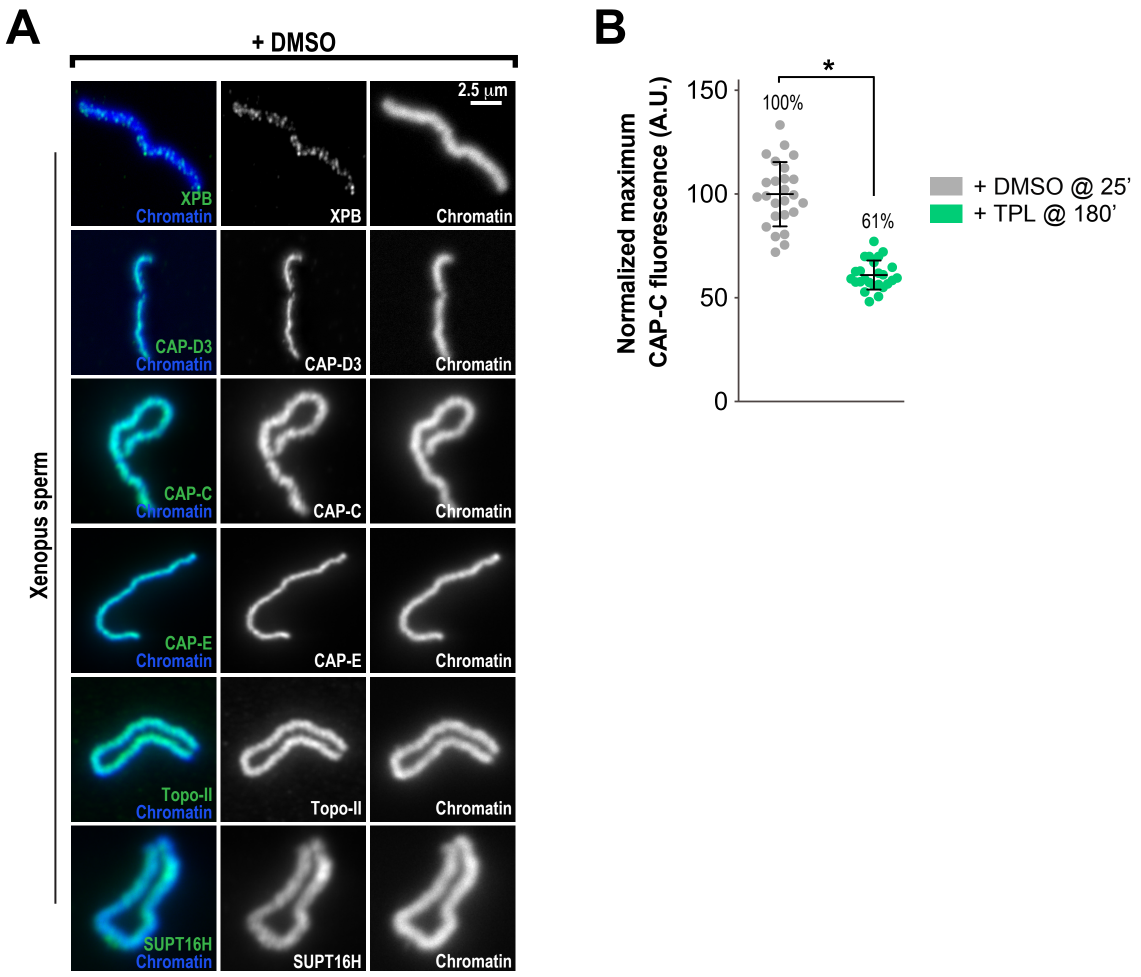


**Figure S3: The TFIIH complex is required for the enrichment of condensins on chromosomes: (A)** Representative immuno-fluorescence images of *Xenopus* sperm nuclei incubated with DMSO or Triptolide treated extracts for 180 min. Chromatids were labelled with Hoechst and anti-XPB, anti-CAP-D3 (condensin II), anti-CAP-C (condensin I & II), anti-CAP-E (condensin I & II), anti-SUPT16H (FACT complex), or anti-Topo II antibodies. Images of individual chromatids are shown. See Figure 3A for images of clusters of chromatids. **(B)** Quantification of fluorescence intensity of CAP-C from experiments in Figures 2B and S2A, normalized to DMSO. n = 50 structures for each condition. Error bars represent SD unless otherwise noted, and asterisks indicate a statistically significant difference (*, $P < 0.001$). A.U., arbitrary units.

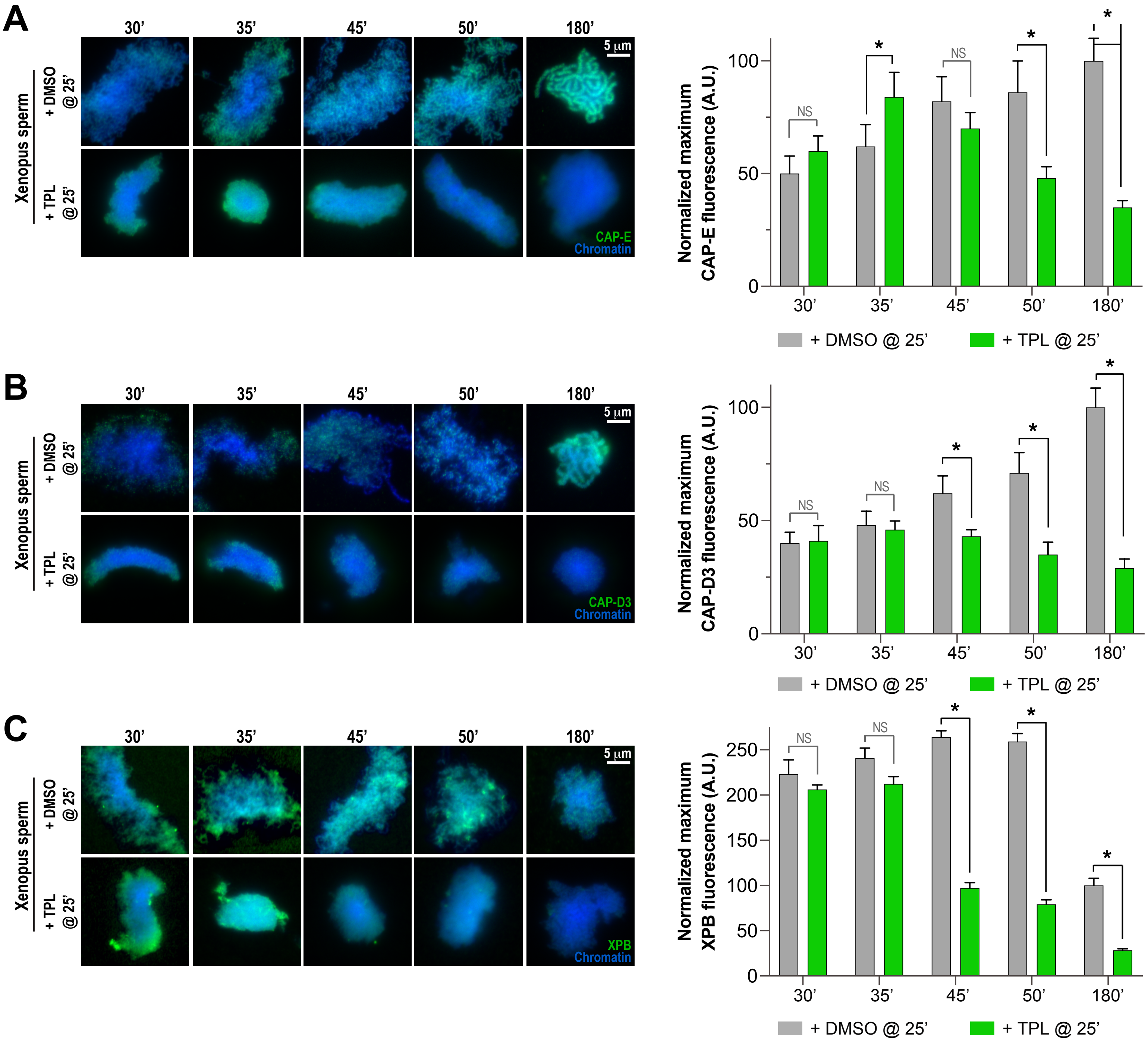


**Figure S4: Triptolide perturbs condensation prior to its effects on condensin levels: (A)** Left: Representative immunofluorescence images of DMSO or triptolide treated extracts at indicated timepoints. Chromatids were labelled with DAPI and anti-CAP-E (condensin I & II). Right: Quantification of fluorescence intensity of CAP-E, normalized to the 180 min DMSO-treated sample. n = 50 structures for each condition. This data is also showin in Figure 4B in gray scale. **(B)** Left: Representative immunofluorescence images of DMSO or triptolide treated extracts at indicated timepoints. Chromatids were labelled with DAPI and anti-CAP-D3 (condensin II). Right: Quantification of fluorescence intensity of CAP-D3, normalized to the 180 min DMSO-treated sample. n = 50 structures for each condition. **(C)** Left: Representative immunofluorescence images of DMSO or triptolide treated extracts at indicated timepoints. Chromatids were labelled with DAPI and anti-XPB (TFIIH complex). Right: Quantification of fluorescence intensity of XPB, normalized to the 180 min DMSO-treated sample. n = 50 structures for each condition.

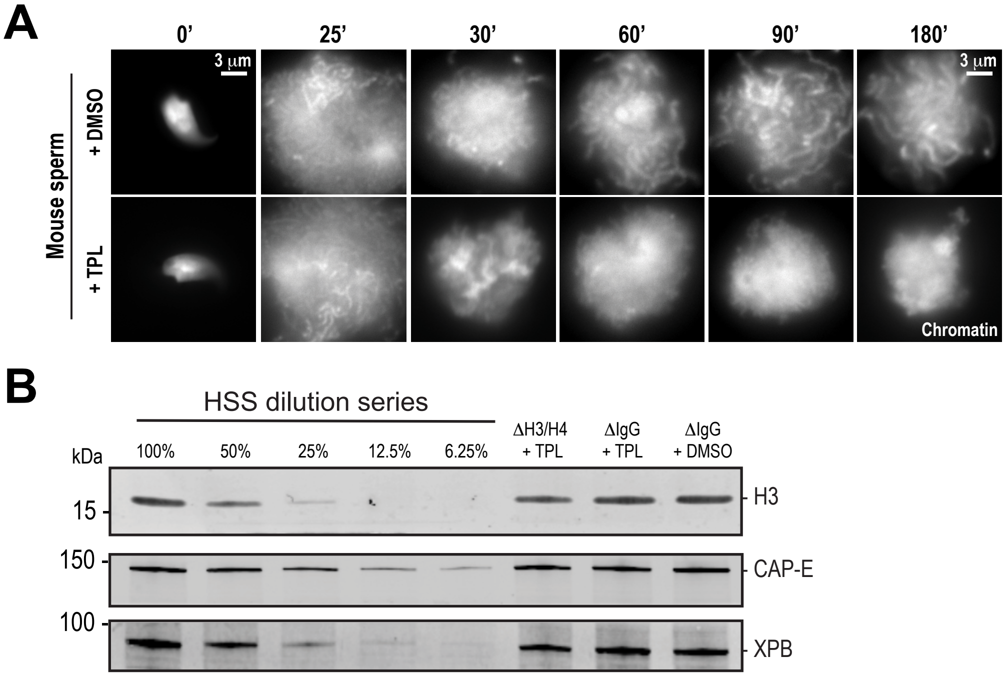


**Figure S5: Effects of triptolide and histone depletion on condensation using mouse sperm: (A)** Representative fluorescence images of chromatid assembly at steady state with mouse sperm nuclei in egg extracts (180 mins after sperm nuclei addition) in the presence of indicated inhibitors. Triptolide or DMSO control was added at 25 mins after nuclei addition. Triptolide was added at 50 μM. **(B)** Western blot for histone H3, CAP-E, and XPB in histone H4K12ac or IgG depleted extracts in the presence of triptolide (TPL) or DMSO.
